# Determining Exon Connectivity in Complex mRNAs by Nanopore Sequencing

**DOI:** 10.1101/019752

**Authors:** Mohan T. Bolisetty, Gopinath Rajadinakaran, Brenton R. Graveley

**Author notes:** These authors contributed equally (Mohan T. Bolisetty, Gopinath Rajadinakaran). Current Address: The Jackson Laboratory for Genomic Medicine, Farmington, CT 06030.

## Abstract

Though powerful, short-read high throughput RNA sequencing is limited in its ability to directly measure exon connectivity in mRNAs containing multiple alternative exons located farther apart than the maximum read lengths. Here, we use the Oxford Nanopore MinION^™^ sequencer to identify 7,899 ‘full-length’ isoforms expressed from four *Drosophila* genes, *Dscam1*, *MRP*, *Mhc*, and *Rdl*. These results demonstrate that nanopore sequencing can be used to deconvolute individual isoforms and that it has the potential to be an important method for comprehensive transcriptome characterization.

## Background

High throughput RNA sequencing has revolutionized genomics and our understanding of the transcriptomes of many organisms. Most eukaryotic genes encode pre-mRNAs that are alternatively spliced (Nilsen and Graveley, 2010). In many genes, alternative splicing occurs at multiple places in the transcribed pre-mRNAs that are often located farther apart than the read lengths of most current high throughput sequencing platforms. As a result, several transcript assembly and quantitation software tools have been developed to address this (Grabherr et al., 2011; Trapnell et al., 2010). While these computational approaches do well with many transcripts, they generally have difficulty assembling and quantitating transcripts that have many isoforms, and for genes with distantly located alternatively spliced regions, they can only infer, and not directly measure, which isoforms may have been present in the original RNA sample (Garber et al., 2011). For example, consider a gene containing two alternatively spliced exons located 2 kbp away from one another in the mRNA. If each exon is observed to be included at a frequency of 50% from short read sequence data, it is impossible to determine whether there are two equally abundant isoforms that each contain or lack both exons, or four equally abundant isoforms that contain both, neither, or only one or the other exon.

Pacific Bioscience sequencing can generate read lengths sufficient to sequence full length cDNA isoforms and several groups have recently reported the use of this approach to characterize the transcriptome (Sharon et al., 2013). However, the large capital expense of this platform can be a prohibitive barrier for some users. Thus, it remains difficult to accurately and directly determine the connectivity of exons within the same transcript. The MinION^™^ nanopore sequencer from Oxford Nanopore requires a small initial financial investment, can generate extremely long reads, and has the potential to revolutionize transcriptome characterization, as well as other areas of genomics.

Several eukaryotic genes can encode hundreds to thousands of isoforms. For example, in *Drosophila*, 47 genes encode over 1,000 isoforms each (Brown et al., 2014). Of these, *Dscam1* is the most extensively alternatively spliced gene known and contains 115 exons, 95 of which are alternatively spliced and organized into four clusters (Schmucker et al., 2000). The exon 4, 6, 9, and 17 clusters contain 12, 48, 33, and 2 exons, respectively. The exons within each cluster are spliced in a mutually exclusive manner and *Dscam1* therefore has the potential to generate 38,016 different mRNA and protein isoforms. The variable exon clusters are also located far from one another in the mRNA and the exons within each cluster are up to 80% identical to one another at the nucleotide level. Together, these characteristics present numerous challenges to characterize exon connectivity within full-length *Dscam1* transcripts for any sequencing platform. Furthermore, though no other gene is as complex as *Dscam1*, many other genes have similar issues that confound the determination of exon connectivity.

We are interested in developing methods to perform simple and robust long-read sequencing of individual isoforms of *Dscam1* and other complex alternatively spliced genes. Here, we use the Oxford Nanopore MinION^™^ to sequence ‘full-length’ cDNAs from four *Drosophila* genes – *Rdl*, *MRP*, *Mhc*, and *Dscam1* – and identify a total of 7,899 distinct isoforms expressed by these four genes.

## Results and Discussion

### Similarity between alternative exons

We were interested in determining the feasibility of using the MinION^™^ nanopore sequencer to characterize the connectivity of distantly located exons in the mRNAs expressed from genes with complex splicing patterns. For the purposes of these experiments, we have focused on four *Drosophila* genes with increasingly complex patterns of alternative splicing (Figure 1). *Resistant to dieldrin* (*Rdl*) contains two clusters each containing two mutually exclusive exons and therefore has the potential to generate 4 different isoforms (Figure 1A). *Multidrug-Resistance like Protein 1* (*MRP*) contains two mutually exclusive exons in cluster 1 and eight mutually exclusive exons in cluster 2, and can generate 16 possible isoforms (Figure 1B). *Myosin heavy chain* (*Mhc*) can potentially generate 180 isoforms due to five clusters of mutually exclusive exons – clusters 1 and 5 contain two exons, clusters 2 and 3 each contain 3 exons, and cluster 4 contains 5 exons. Finally, *Dscam1* contains 12 exon 4 variants, 48 exon 6 variants, 33 exon 9 variants (Figure 1D) and 2 exon 17 variants (not shown) and can potentially express 38,016 isoforms. For this study, however, we have focused only on the exon 3 through exon 10 region of *Dscam1*, which encompasses the 93 exon 4, 6, and 9 variants, and 19,008 potential isoforms (Figure 1D).

**Figure 1.**
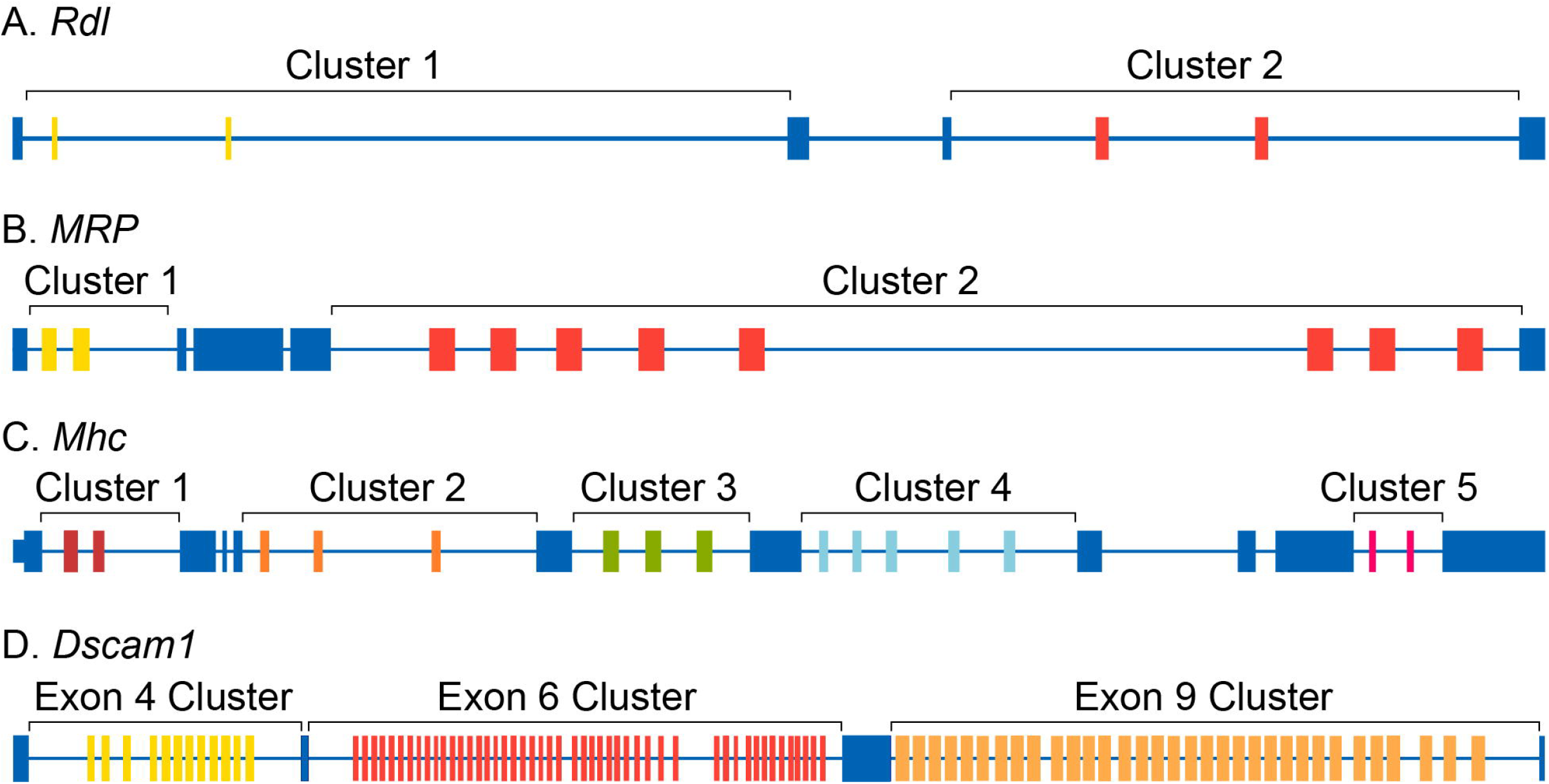
Schematic of the exon-intron structures of the genes examined in this study. **A.** The *Rdl* gene contains two clusters (cluster one and two) which each contain two mutually exclusive exons. **B.** The *MRP* gene contains contains two and eight mutually exclusive exons in clusters one and two, respectively. **C.** *Mhc* contains two mutually exclusive exons in clusters one and five, three mutually exclusive exons in clusters two and three, and five mutually exclusive exons in cluster four. **D.** The *Dscam1* gene contains 12, 48, and 33 mutually exclusive exons in the exon four, six and nine clusters, respectively. For each gene, the constitutive exons are colored blue, while the variable exons are colored yellow, red, orange, green or light blue.

Because our nanopore sequence analysis pipeline uses LAST to perform alignments (Frith et al., 2010), we aligned all of the *Rdl*, *MRP*, *Mhc*, and *Dscam1* exons within each cluster to one another using LAST to determine the extent of discrimination needed to accurately assign nanopore reads to a specific exon variant. For *Rdl*, each variable exon was only aligned to itself, and not to the other exon in the same cluster (data not shown). For *MRP*, the two exons within cluster 1 only align to themselves, and though the 8 variable exons in cluster 2 do align to other exons, there is sufficient specificity to accurately assign nanopore reads to individual exons (Figure 2a). For *Mhc*, the variable exons in the cluster 1 and cluster 5 do not align to other exons, and the variable exons in cluster 2, cluster 3 and cluster 4 again align with sufficient discrimination to identify the precise exon present in the nanopore reads (Figure 2b). Finally, for *Dscam1*, the difference in the LAST alignment scores between the best alignment (each exon to itself) and the second, third and fourth best alignments are sufficient to identify the *Dscam1* exon variant (Figure 2c). This analysis indicates that for each gene in this study, LAST alignment scores are sufficiently distinct to identify the variable exons present in each nanopore read.

**Figure 2.**
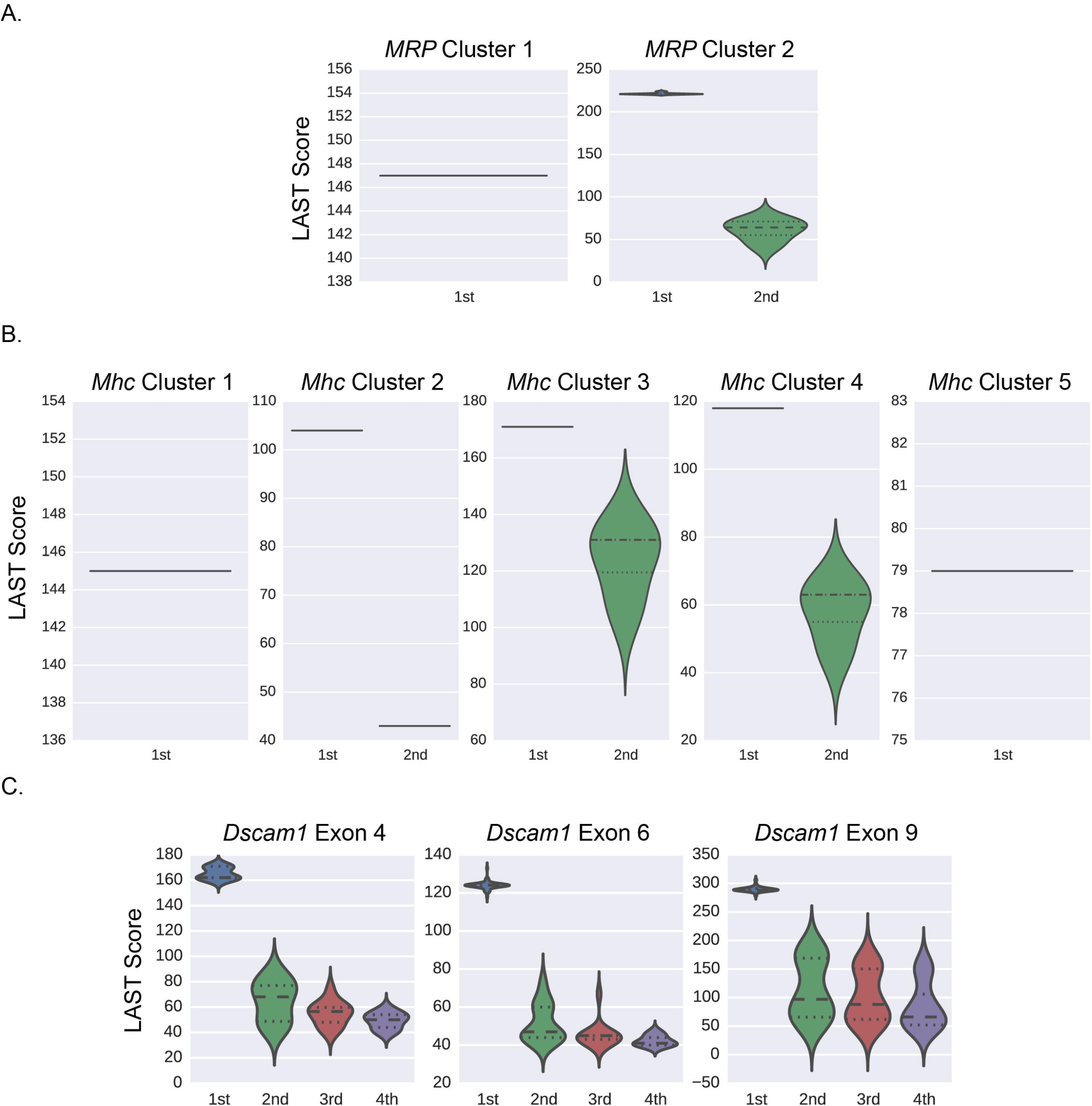
Similarity distance between the variable alternative exons of *MRP*, *Mhc*, and *Dscam1*. **A.** Schematic of the organization of the exon 3 through exon 10 region of *Dscam1*. **B.** Violin plots of the LAST alignment scores of each variable exon within each cluster to themselves (1st), and to the exons with the second (2nd), third (3rd) and fourth (4th) best alignments.

### Optimizing Template Switching in Dscam1 cDNA libraries

Template switching can occur frequently when libraries are prepared by PCR and can confound the interpretation of results (McManus et al., 2010; Plocik and Graveley, 2013). For example, CAM-Seq (Sun et al., 2013) and a similar method we independently developed called Triple-Read sequencing (Roy et al., 2015) to characterize *Dscam1* isoforms, were found to have excessive template switching due to amplification during the library prep protocols. To assess template switching in our current study, we generated a spike-in mixture of in vitro transcribed RNAs representing six unique *Dscam1* isoforms – *Dscam1*^*4.2,6.32,9.31*^, *Dscam1*^*4.1,6.46,9.30*^, *Dscam1*^*4.3,6.33,9.9*^, *Dscam1*^*4.12,6.44,9.32*^, *Dscam1*^*4.7,6.8,9.15*^, and *Dscam1*^*4.5,6.4,9.4*^. We used 10 pg of this control spike-in mixture and prepared libraries for MinION^™^ sequencing by amplifying the exon 3 through exon 10 region for 20, 25 or 30 cycles of RT-PCR. We then end-repaired and dA-tailed the fragments, ligated adapters, and sequenced the samples on a MinION^™^ (7.3) for 12 hours each. We obtained 33,736, 8,961 and 7,511 base-called reads from the 20, 25, and 30 cycle libraries, respectively. Consistent with the size of the exon 3 to 10 cDNA fragment being 1,806 −1,860 bp in length, depending on the precise combination of exons it contains, most reads we observed were in this size range (Figure 3A). We used Poretools (Loman and Quinlan, 2014) to convert the raw output files into fasta format and then used LAST to align the reads to a LAST database containing each variable exon. From these alignments, we identified reads that mapped to all three exon clusters, as well as the exon with the best alignment score within each cluster. When examining the alignments to each cluster independently, we found that for these spike-in libraries, all reads mapped uniquely to the exons present in the input isoforms. Therefore, any observed isoforms that were not present in the input pool were a result of template-switching during the RT-PCR and library prep protocol and not due to false alignments or sequencing errors.

**Figure 3.**
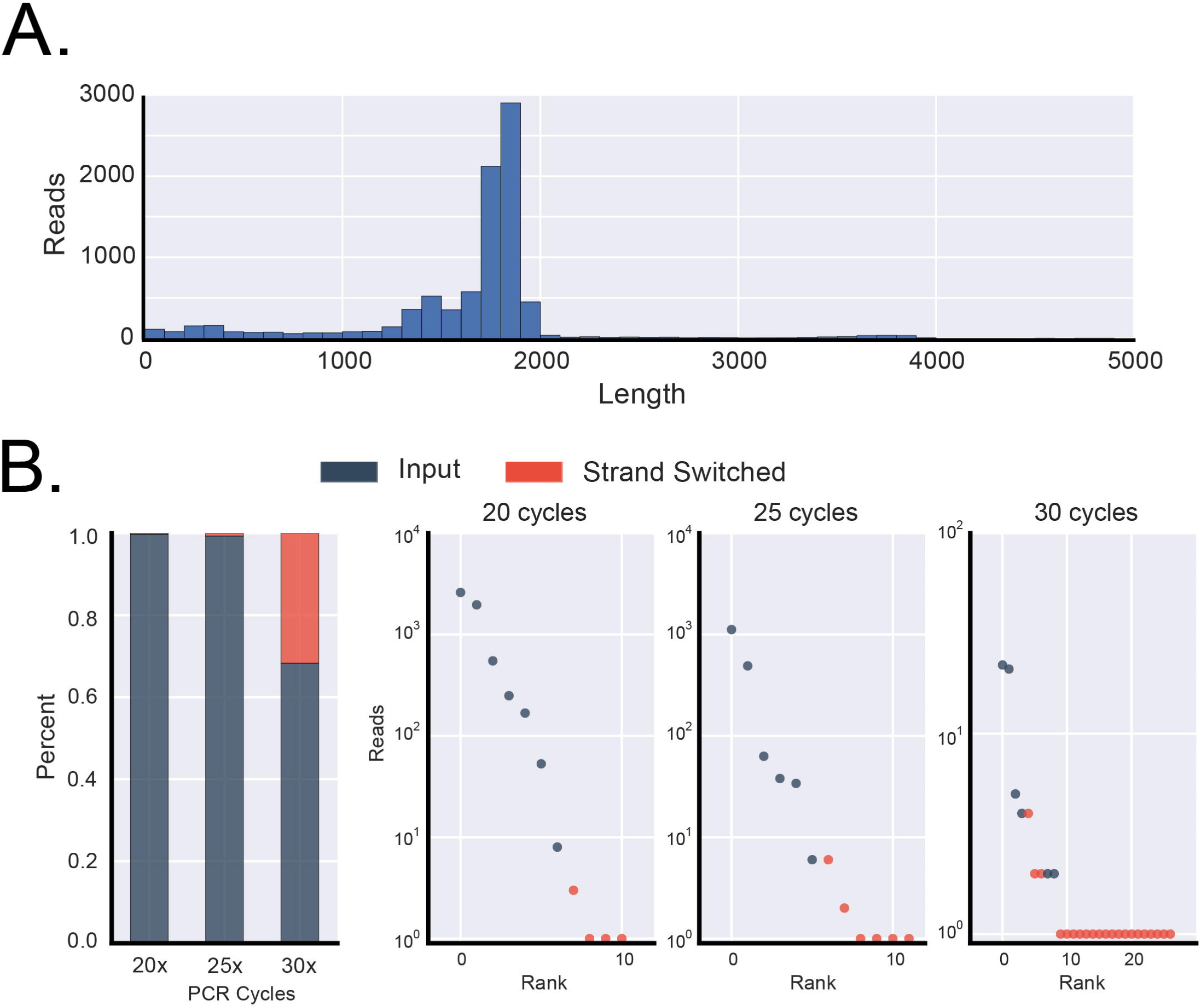
Optimized RT-PCR minimizes template-switching for MinION sequencing. **A.** Histogram of read lengths from MinION sequencing of *Dscam1* spike-ins from the library generated using 25 cycles of PCR. **B.** Bar plot indicating the extent of template-switching in *Dscam1* spike-ins at different PCR cycles. The blue portions indicate the fraction of reads corresponding to input isoforms while the red portions correspond to the fraction of reads corresponding to template-switched isoforms. **C.** Plots of the rank order versus number of reads (log10) for the 20, 25 and 30 cycle libraries. The blue dots indicate input isoforms while the red portions correspond to template-switched isoforms.

When comparing the combinations of exons within each read to the input isoforms, we observed that 32% of the reads from the 30 cycle library corresponded to isoforms generated by template switching (Figure 3B). The template switched isoforms observed by the greatest number of reads in the 30 cycle library were due to template switching between the two most frequently sequenced input isoforms. In most cases, template switching occurred somewhere within exon 7 or 8 and resulted in a change in exon 9. However, the extent of template switching was reduced to only 1% in the libraries prepared using 25 cycles, and to 0.2% in the libraries prepared using 20 cycles of PCR (Figure 3B). Again, for these two libraries the most frequently sequenced template switched isoforms involved the input isoforms that were also the most frequently sequenced. These experiments demonstrate that the MinION^™^ nanopore sequencer can be used to sequence ‘full length’ *Dscam1* cDNAs with sufficient accuracy to identify isoforms and that the cDNA libraries can be prepared in a manner that results in a very small amount of template-switching.

### Dscam1 isoforms observed in adult heads

To explore the diversity of *Dscam1* isoforms expressed in a biological sample, we prepared a *Dscam1* library from RNA isolated from *D. melanogaster* heads prepared from mixed male and female adults using 25 cycles of PCR and sequenced it for 12 h on the MinION^™^ nanopore sequencer obtaining a total of 159,948 reads of which 78,097 were template reads, 48,474 were complement reads and 33,377 were 2D reads (Figure 4A). We aligned the reads individually to the exon 4, 6, and 9 variants using LAST. A total of 28,971 reads could be uniquely or preferentially aligned to a single variant in all three clusters. For further analysis, we used all 2D read alignments (16,419) and only 1D reads when both template and complement aligned to same variant exons (31). We observed 92 of the 93 potential exon 4, 6 or 9 variants – only exon 6.11 was not observed in any read (Figure 4D). Over their entire lengths, the 2D reads that map specifically to one exon 4, 6 and 9 variants map with an average 90.37% identity and an average LAST score of ∼1200 (Supplemental Figure 1). The 16,450 full length reads correspond to 7,874 unique isoforms, or 42% of the 18,612 possible isoforms given the exon 4, 6 and 9 variants observed. We note, however, that while 4,385 isoforms were represented by more than one read, 3,516 of isoforms were represented by only 1 read indicating that the depth of sequencing has not reached saturation (Figure 4B and 4C). The most frequently observed isoforms were *Dscam1*^*4.1,6.12,9.30*^ and *Dscam1*^*4.1,6.1,9.30*^ which were observed with 35 and 25 reads respectively (Figure 4D).

**Figure 4.**
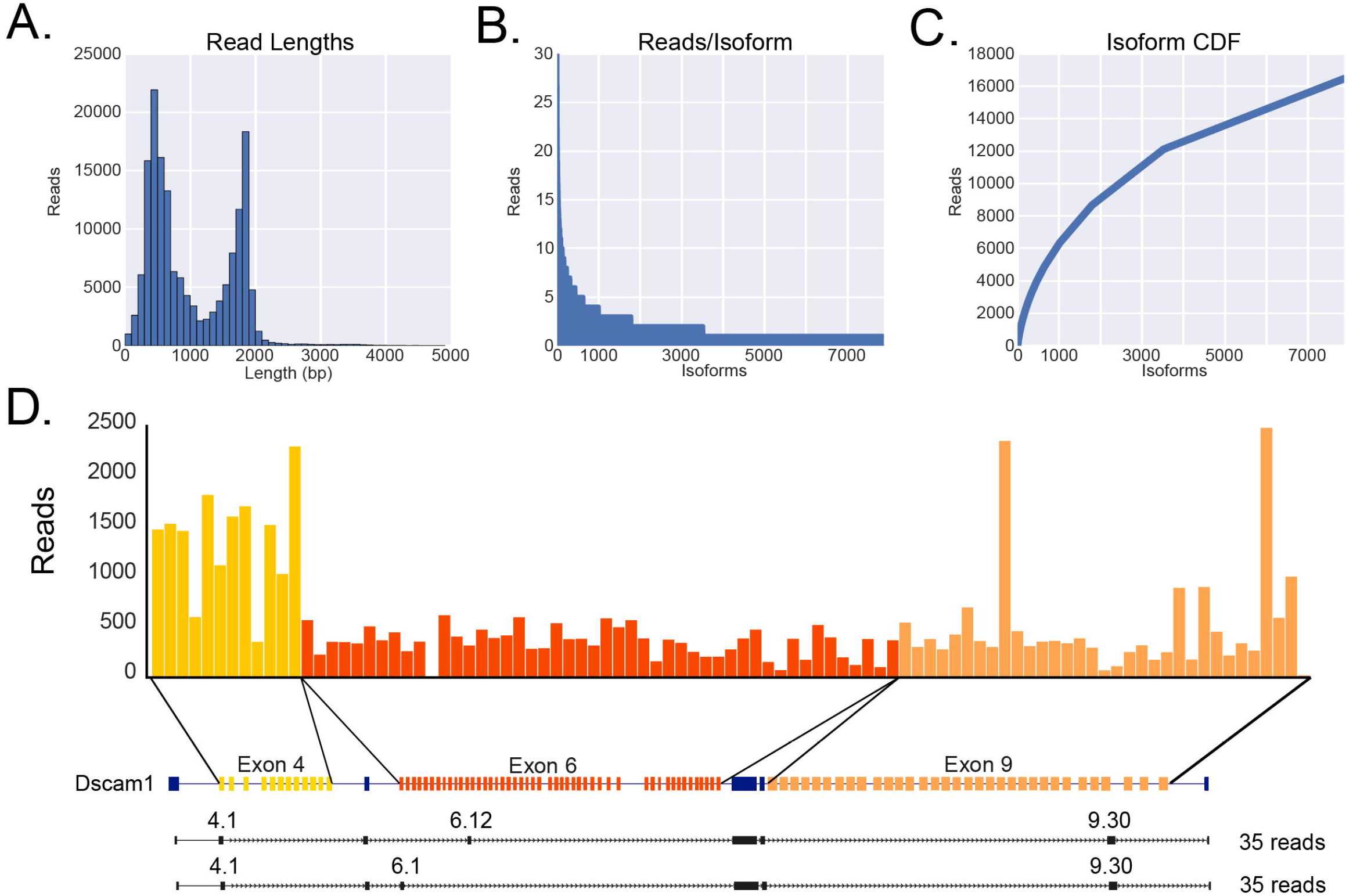
MinION sequencing of *Dscam1* identifies 6,516 isoforms. **A.** Histogram of read length distribution for Drosophila head samples. **B.** The total number of *Dscam1* isoforms identified from MinION sequencing. **C.** Cumulative distribution of *Dscam1* isoforms with respect to expression. **D.** Deconvoluted expression of *Dscam1* exon cluster variants. **E.** Isoform connectivity of two highly expressed *Dscam1* isoforms.

### Nanopore sequencing of ‘full-length’ Rdl, MRP, and Mhc isoforms

To extend this approach to other genes with complex splicing patterns, we focused on *Rdl*, *MRP*, and *Mhc* which have the potential to generate 4, 16, and 180 isoforms, respectively. We prepared libraries for each of these genes by RT-PCR using primers in the constitutive exons flanking the most distal alternative exons using 20 cycles of PCR, pooled the three libraries and sequenced them together on the MinION^™^ nanopore sequencer for 12 hours obtaining a total of 22,962 reads. The input libraries for *Rdl*, *MRP* and *Mhc* were 567 bp, 1,769-1,772 bp, and 3,824 bp, respectively. The raw reads were aligned independently to LAST indexes of each cluster of variable exons. The alignment results were then used to assign reads to their respective libraries, identify reads that mapped to all variable exon clusters for each gene, and the exon with the best alignment score within each cluster (Figure 5). In total, we obtained 301, 337 and 112 full length reads for *Rdl*, *MRP*, and *Mhc*, respectively. From these reads, we observed a total of 4 *Rdl* isoforms, 9 *MRP* isoforms and 12 *Mhc* isoforms.

**Figure 5.**
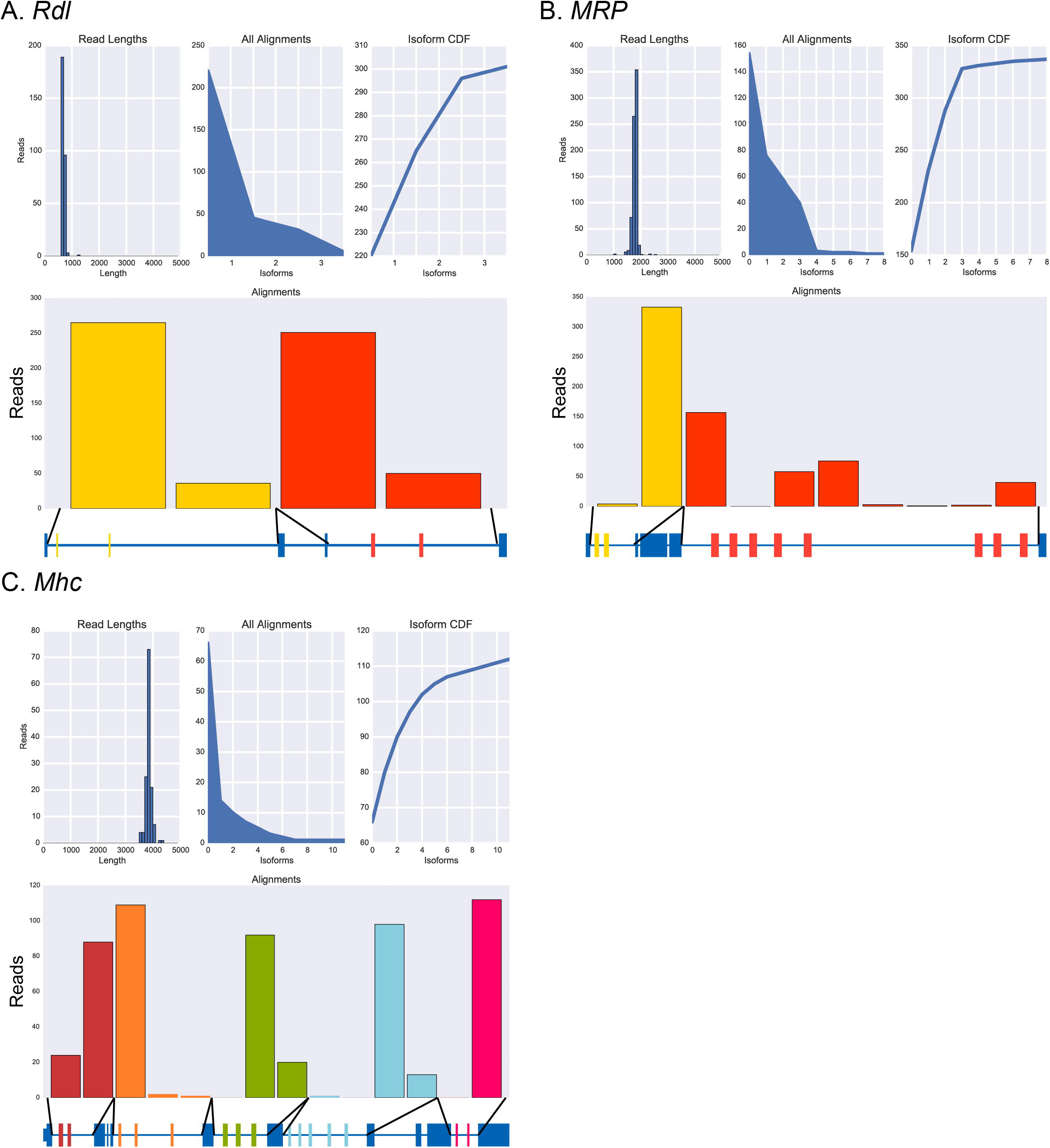
MinION sequencing of *Rdl*, *MRP* and *Mhc* Isoforms. Histogram of read lengths, number of reads per isoform, cumulative distribution of isoforms with respect to expression, and number of reads per alternative exon for *Rdl* (**A**), *MRP* (**B**), and *Mhc* (**C**).

## Conclusions

Here we have demonstrated that nanopore sequencing can be used to easily determine the connectivity of exons in a single transcript, even for the most complicated alternatively spliced genes. This is an important advance because short-read sequence data cannot be used to conclusively determine which exons are present in the same RNA molecule, especially for complex alternatively spliced genes. We anticipate that nanopore sequencing of whole transcriptomes, rather than targeted genes as we have performed here, will be a rapid and powerful approach for characterizing isoforms, especially with improvements in the throughput and accuracy of the technology, and the simplification and/or elimination of the time-consuming library preparations.

## Materials and Methods

### Drosophila strains

*Drosophila melanogaster y*; *cn b sp* (stock: 2057, Bloomington) were maintained and raised at room temperature.

### Spike-in preparation

Total RNA from about 30 heads was extracted using Trizol reagent. 1 μg of total RNA was used to synthesize cDNA using random hexamers with SuperScript II (Invitrogen, Cat No: 18064) in a 20 μl reaction. 2 μl of cDNA reaction was used to amplify *Dscam1* exons 4 through 9 using the primers exon 3 and exon 10 with LongAmp (New England Biolabs, Cat No: M0323) in a 50 μl reaction volume with the following PCR condition: initial denaturation at 94°C for 30S, denaturation at 94°C for 15S, annealing at 58°C for 15S, extension at 65°C for 100S (40X cycle), final extension at 65°C for 10 mins. The PCR amplicons were purified using MinElute PCR purification kit (Qiagen) and eluted in 20 μl ultrapure water. The eluted amplicons were then cloned into a vector with both T7 and SP6 dual promoters (Life Technologies, Cat No: K4600) and transformed into Top10 shot cells. A total of 96 colonies were sequenced to identify exon variant sequences in individual clones. Six individual colonies containing a single, non-overlapping, unique exon variants were used to make spike-in RNAs. The vector containing the *Dscam1* insert and the T7, SP6 promoter sequences were amplified using M13F and M13R primers. The SP6 oriented clones were individually amplified using T7 overhang primers to facilitate *in vitro* transcription of all clones from T7 promoter using transcription kit. Following transcription, 1 μl RNA (1 μg/μl) of each of the six clones were mixed and a 10 fold serial dilution was made with concentration ranging from 100 ng/μl to 1 pg/μl. cDNA was synthesized using SuperScript II (Invitrogen, Cat No: 18064) and a 2.5 μl cDNA from 10 pg/μl reaction was used in the 25 μl Phusion PCR with the following conditions: initial denaturation at 95°C for 30S, denaturation at 95°C for 10S, annealing at 64.7°C for 12S, extension at 72°C for 40S (20X, 25X, and 30X cycles), final extension at 72°C for 5 mins, using primers CGGATCCATTATCTCCCGGGACG (*Dscam1* exon 3) and CGGATCCCTGGGCGAAGGCC (*Dscam1* exon 10 reverse).

### Amplicon library preparation and Oxford Nanopore Sequencing

The library preparation for amplicon sequencing was done using SQK-MAP003 following manufacturers protocol (ONT). Briefly, a total of 850 ng (spike-in) and 1 μg (mix heads) in 80 μl was end repaired using NEBNext End Repair Module (New England Biolabs, Cat No: E6050) and followed by dA tailing using NEBNext dA Tailing Module (New England Biolabs, Cat No: E6053). The dA tailed amplicons were then adapter ligated in a total of 100 μl reaction volume and incubated at room temperature for 10 mins. This reaction mixture was then purified using Agencourt AMPure XP (Beckman Coulter Inc., cat. no. A63880) beads and washed and eluted in Nanopore supplied reagents in 25 μl ultrapure water. This pre-sequencing mix was added with the fuel mix and EP buffer and loaded on the R7.3 flow cell and sequenced.

### Data analysis

Poretools (version 0.3.0) was used to extract fasta reads from Basecalled fast5 files. Exon cluster specific LAST indices were made using lastdb with default parameters. The reads were then aligned using lastal independently to these LAST indices using following parameters: -s 2 -T 0 -Q 0 -a 1. Reads that aligned to all 3 clusters were parsed from all alignments and used for further processing. The top scoring alignment was used for reads that aligned to multiple variants. iPython notebooks containing all the analysis and code are available at github/mohanbolisetty/dscam_nanopore. MAF files from LAST alignments were converted to SAM or PSL formats using maf-convert.py.

### Accession codes

Sequence reads are currently being submitted to a public archive and will be made available as soon as possible.

## ACKNOWLEDGMENTS

We thank members of the Graveley laboratory for comments on this work and the Oxford Nanopore for reagents, MinIONs, and the opportunity to participate in the Oxford Nanopore MinION Access Programme (MAP). This work was supported in part by US National Institutes of Health grant R01GM067842 and the John and Donna Krenicki Endowment Fund to B.G. M.B was funded by AHA founders affiliate postdoctoral fellowship grant 14POST18750000.

### AUTHOR CONTRIBUTIONS

M.B., G.R., and B.G. conceived of the idea and designed the experiments. M.B. and G.R. performed the experiments. M.B. and B.G. conducted bioinformatic analyses and wrote the paper with input from G.R.

### COMPETING FINANCIAL INTERESTS

The authors declare no competing financial interests.

---

**Supplementary Figure 1. Percent identities of *Dscam1* reads from MinION sequencer. A.**Percent identities of MinION reads mapping to individual exon variants in 4, 6, and 9 clusters. **B.** Percent identities of alignments with respect to template, complement and two directions (sequencing both template and complements).

## References

Brown JB, Boley N, Eisman R, et al. Diversity and dynamics of the Drosophila transcriptome. Nature. 2014; 512(7515):393–399. doi:10.1038/nature12962.

Frith MC, Hamada M, Horton P. Parameters for accurate genome alignment. BMC Bioinformatics 2010, 11:80. doi:10.1186/1471-2105-11-80.

Garber, M., Grabherr, M.G., Guttman, M., and Trapnell, C. (2011). Computational methods for transcriptome annotation and quantification using RNA-seq. Nat Meth 8, 469–477.

Grabherr, M.G., Haas, B.J., Yassour, M., Levin, J.Z., Thompson, D.A., Amit, I., Adiconis, X., Fan, L., Raychowdhury, R., Zeng, Q., et al. (2011). Full-length transcriptome assembly from RNA-Seq data without a reference genome. Nat. Biotechnol. 29, 644–652.

Loman NJ, Quinlan AR. Poretools: a toolkit for analyzing nanopore sequence data. Bioinformatics. 2014;30(23):3399-3401. doi:10.1093/bioinformatics/btu555.

McManus, C.J., Duff, M.O., Eipper-Mains, J., and Graveley, B.R. (2010). Global analysis of trans-splicing in Drosophila. Proc Natl Acad Sci USA 107, 12975–12979.

Nilsen, T.W., and Graveley, B.R. (2010). Expansion of the eukaryotic proteome by alternative splicing. Nature 463, 457–463.

Plocik, A.M., and Graveley, B.R. (2013). New Insights from Existing Sequence Data: Generating Breakthroughs without a Pipette. Mol Cell 49, 605–617.

Roy CK, Olson S, Graveley BR, Zamore PD, Moore MJ. Assessing long-distance RNA sequence connectivity via RNA-templated DNA-DNA ligation. eLife. 2015 Apr 13; 4. doi:10.7554/eLife.03700.

Schmucker, D., Clemens, J.C., Shu, H., Worby, C.A., Xiao, J., Muda, M., Dixon, J.E., and Zipursky, S.L. (2000). Drosophila Dscam is an axon guidance receptor exhibiting extraordinary molecular diversity. Cell 101, 671– 684.

Sharon D, Tilgner H, Grubert F, Snyder M. A single-molecule long-read survey of the human transcriptome. Nature Biotechnology 2013, 31:1009–1014. doi:10.1038/nbt.2705.

Sun W, You X, Gogol-Döring A, et al. (2013) Ultra-deep profiling of alternatively spliced Drosophila Dscam isoforms by circularization-assisted multi-segment sequencing. The EMBO Journal. 32:2029–2038.

Trapnell, C., Williams, B.A., Pertea, G., Mortazavi, A., Kwan, G., van Baren, M.J., Salzberg, S.L., Wold, B.J., and Pachter, L. (2010). Transcript assembly and quantification by RNA-Seq reveals unannotated transcripts and isoform switching during cell differentiation. Nat. Biotechnol. 28, 511–515.

